# A temporal hierarchy determines epigenetic aging

**DOI:** 10.1101/2025.08.13.669476

**Authors:** Matteo Ciarchi, Benjamin D. Simons, Steffen Rulands

## Abstract

Aging involves processes spanning orders of magnitude in time, from fast events that occur at the molecular scale to the slow decrease of physiological function. Whether and how fast molecular events lead to the slow progression of aging, and what ultimately sets the timescale of aging, is not understood. Here, by focusing on dynamic changes in DNA methylation, we show how aging phenomena on long timescales emerge from the kinetics of fast molecular processes, providing a bridge between temporal scales. By combining DNA methylation sequencing data across a range of timescales with a statistical modeling-based approach, we show that DNA methylation aging is governed by a three-fold hierarchy of processes that dominate on distinct timescales: individual stochastic events in which enzymes interact with the DNA and with each other (milliseconds); the convergence of molecular concentrations to steady states (days to months); and stochastic transitions between these steady states (years to decades). Our findings provide a unified picture of how DNA methylation aging arises across temporal scales.

---

Aging is manifest in multiple processes on vastly different timescales, from protein binding to DNA that occurs on the millisecond timescale, to the organism-level decay of biological function that occurs in humans on the timescale of decades (Fig. 1a) (1). At the molecular scale, alterations of epigenetic modifications of DNA and chromatin are closely associated with aging, including DNA methylation, histone modifications, and chromatin remodeling (2; 3; 4; 5). Among these, the primary layer of epigenetic modification is DNA methylation, which affects cytosines in a CpG context. During development, *de novo* DNA methylation is established through the action of the methyltransferase enzymes, DNMT3A and DNMT3B (6). DNA replication leads to the creation of hemimethylated sites, which are subsequently corrected by the action of a complex of enzymes involving DNMT1 and UHRF1 (7; 8). In the absence of DNMT1, DNA methylation levels can become passively diluted over several rounds of cell division (9). DNA methylation marks can also be actively removed by the action of TET1/2/3 enzymes and the base-pair excision and repair pathway (10).

**Figure 1.**
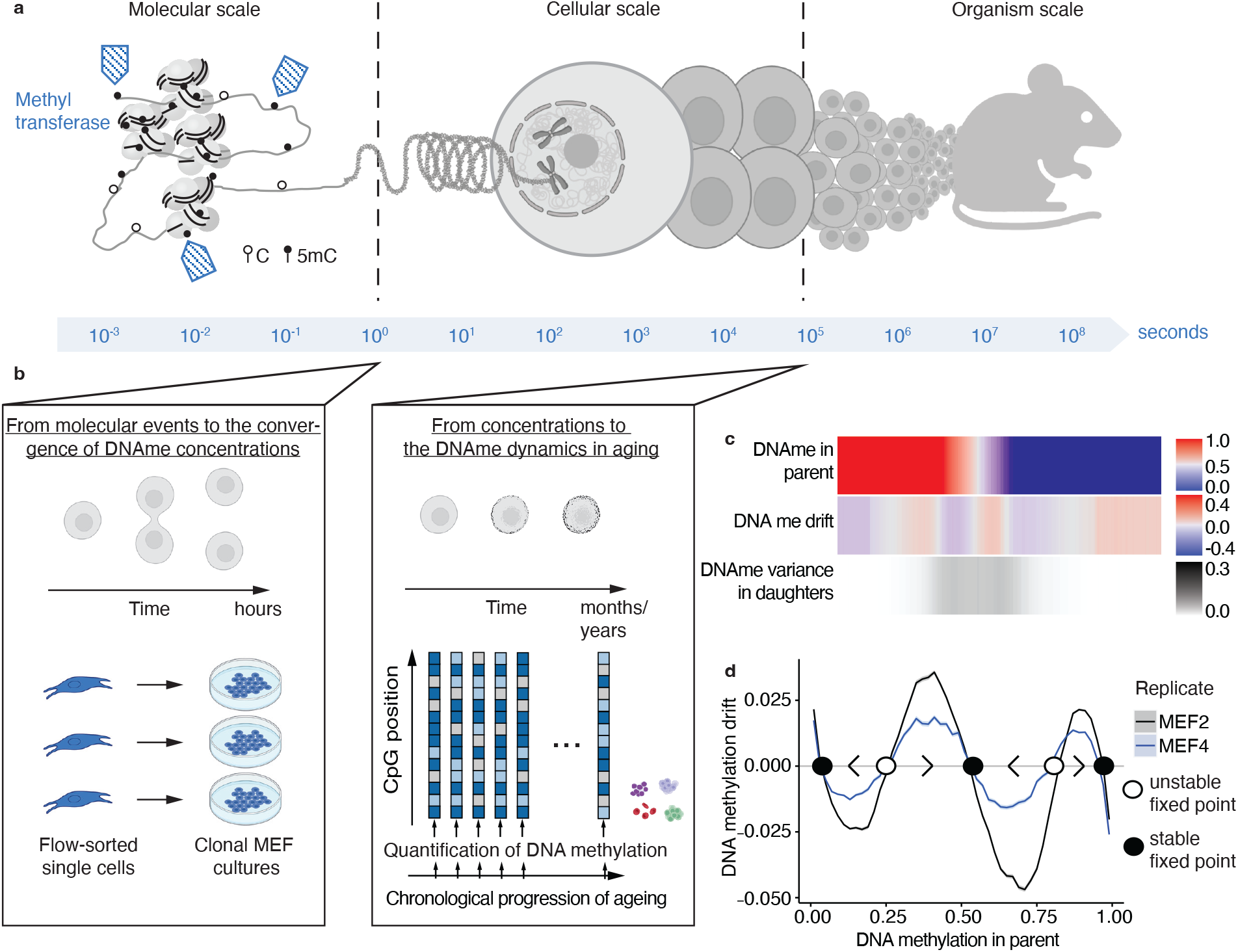
Spatial and temporal scales in aging. **a** Schematic showing the range of temporal scales contributing to aging.**b** Schematic showing the two different experimental approaches used to generate the data used in this study and how they relate to the temporal scales of aging. On short timescales, we used clonal DNA methylation data of MEF cells (left). For DNA methylation dynamics on longer timescales, we drew on aging experiments measuring DNA methylation over different ages and different tissues. **c** Heatmap showing the DNA methylation drift, parental DNA methylation, and DNA methylation variance over daughter cells for the clonal MEF data (27). (For details, see main text.) **d** DNA methylation drift as a function of parent DNA methylation for the two cell lines of the clonal MEF data. Circles (open and filled) indicate the position of dynamical fixed points and their stability. Arrows indicate the direction in which DNA methylation levels tend to change over time.

DNA methylation, as well as histone modifications, have been found to change systematically during aging (11; 12). Indeed, statistical and machine learning models have been shown to accurately predict chronological age across tissues and species in bulk (13) and at the single-cell level (14; 15; 16; 17; 18; 19). These “DNA methylation clock” models partially measure the accumulation of stochastic variation in DNA methylation over time, while others have emphasized contributions of deterministic dynamics (20; 21; 22). In line with a stochastic time-evolution is the observation that average DNA methylation levels tend to converge to intermediary values (23). Gene-related CpG islands, which tend to be lowly methylated, gain methylation over time. Conversely, DNA methylation tends to decrease in genomic regions associated with heterochromatin, such as repetitive elements and transposons, and regions of low CpG density, which are highly methylated under normal conditions (24). Because of its tight association with biological age, DNA methylation is a suitable framework for understanding how molecular events at the millisecond scale lead to the slow progression of aging on time scales that are orders of magnitude slower.

How then do the molecular processes that control DNA methylation alterations at the millisecond timescale integrate in the long-term dynamics of DNA methylation during aging? Conceptually, there are two possible answers to this question: First, DNA methylation aging could be driven by a slow process that is extrinsic to the dynamics of DNA methylation. For example, the production of methyl groups for DNA methylation is coupled to metabolism via one-carbon and polyamine metabolism (25). Slow changes in metabolism during aging could therefore lead to temporal changes in the localization of DNA methylation marks with time. Alternatively, DNA methylation aging could be driven intrinsically as a result of the accumulation of fast molecular processes altering the epigenome (26).

Here, combining experiments on different temporal scales and a dynamical systems approach, we link the fast molecular processes that establish DNA methylation to the epigenetic processes occurring during cell division, and ultimately, the much slower dynamics of DNA methylation during aging. Our results show that DNA methylation aging is the result of the accumulation of these fast molecular processes. Specifically, drawing on experiments that address the short-term dynamics of DNA methylation, we derive a minimal biophysical model that links individual molecular events to the time evolution of DNA methylation over several rounds of cell divisions. In this model, the genome-wide DNA methylation dynamics is governed by a single parameter reflecting the relative strength of local coordination and stochasticity. We then use longterm longitudinal experiments quantifying DNA methylation during aging to link these processes to DNA methylation drift on the timescales of aging.

## Results

To investigate how the fast molecular events regulating methylation of individual CpGs are related to DNA methylation aging dynamics on much slower timescales, we made use of two complementary experimental designs: the first focused on short-term dynamics over a few rounds of cell divisions and the second on the long-term dynamics of aging (Fig. 1b). To quantify short-term DNA methylation dynamics, we used previously published target-capture bisulfite sequencing data, where mouse embryonic fibroblast cells (MEFs) collected at embryonic day (E)13.5 were immortalized to produce two different cell lines (27). From each line, 7 cultures were prepared from single cells, and clones were subject to high-throughput target-capture bulk bisulfite sequencing. To study DNA methylation dynamics over the long term, we used longitudinal reduced-representation bisulfite sequencing (RRBS-seq) studies in a range of tissues (28), as well as a single-cell bisulfite sequencing experiment from mouse blood (29). To bridge temporal scales, in the first step, we made use of the data from the clonal MEF lines to relate individual molecular events to the convergence of molecular concentrations to their respective steady states. Then, in the second step, we made use of the longitudinal data on aging to extrapolate these findings to the timescale of aging.

### From molecular events to DNA methylation drift

To relate the molecular kinetics of DNA methylation turnover to the time evolution of DNA methylation concentrations toward their steady state, we first quantified how DNA methylation levels evolve from a given starting value in the parent to the clonal daughter populations. In bisulfite sequencing experiments, the quantification of DNA methylation takes binary values in single-cell sequencing or continuous values between 0 and 1 (termed beta values) for bulk sequencing. Since we consider both types of experiments in this work, we collectively refer to the experimental readout as a DNA methylation level. To quantify the change in DNA methylation, for each biological replicate and CpG site, we determined the DNA methylation drift, Δ*m*, defined as the difference between the DNA methylation level averaged over daughter clones and the corresponding value in the parent colony. At the same time, we computed the variance in DNA methylation levels over the seven clonal samples. As expected, the variance across the clonal cell lines was highest for CpGs with intermediary average levels of DNA methylation (Fig. 1c) (27). The data also showed a non-monotonic relation between the DNA methylation level of a given CpG in the parent population and the DNA methylation drift (Fig. 1c).

To understand the origin of this non-monotonicity, we then grouped CpG sites into bins of similar DNA methylation levels in the parent population and calculated the average DNA methylation drift for each bin. The average DNA methylation drift showed a highly nonlinear and non-monotonic dependence on the parent DNA methylation level (Fig. 1d). By dividing the DNA methylation drift by the difference in time, Δ*t*, Fig. 1d can be interpreted as an effective DNA methylation-dependent “force”, *f* (*m*), indicating the direction in which DNA methylation evolves over time. We can, therefore, adopt a dynamical systems perspective in which the time-evolution of DNA methylation is defined by two properties: first, the fixed points, i.e., the DNA methylation levels at which the drift vanishes, *f* (*m*) = 0; and second, the stability of these fixed points, as determined by the gradient of *f* (*m*) evaluated at the fixed points. Specifically, a fixed point is stable if the dynamics in its vicinity converge towards the fixed point such that 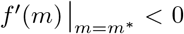, where *m*^∗^ is the value of the fixed point. By contrast, the dynamics near unstable fixed points evolve away from these fixed points, such that 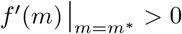. With these definitions, we find that the whole-genome aggregate DNA methylation dynamics is governed by five fixed points, three of which (*m*^∗^ = 0, *m*^∗^ ≈ 1*/*2, and *m*^∗^ = 1) are stable, while two (*m* ≈ 1*/*4 and *m* ≈ 3*/*4) are unstable (Fig. 1d).

To link individual molecular events to DNA methylation dynamics on the timescale of aging, we first needed to show that the observed behavior of the DNA methylation drift is caused by molecular events. To this end, we reasoned that the observed systematic variation in the drift must result from nonlinear feedback. In bulk sequencing, such as target-capture bisulfite sequencing, the accumulated reads reflect an aggregation of strand-, allele-, and cell-specific reads. In principle, the nonlinear feedback governing the DNA methylation dynamics could, therefore, originate on the molecular, cellular, or population scale.

At the scale of the cell population, cell proliferation is the predominant process driving DNA methylation changes in MEF cultures. Here, the copying of hemimethylated sites to the newly-replicated DNA strand following cell division can lead to DNA methylation drift (Fig. 2a). During DNA replication, a complex of proteins, including DNMT1, detects hemimethylated sites and copies DNA methylation marks onto the newly-replicated DNA strand. The processive action of DNMT1 enzymes leads to genomic correlations of this process (30). Indeed, the presence of unstable fixed points at DNA methylation values of around 1*/*4 and 3*/*4 could derive from the correction of hemimethylated sites during DNA replication. However, while this mechanism can explain the drift towards stable fixed points at 0, 1*/*2 and 1 on the cellular scale, it cannot explain the nonlinear drift on the population scale, as observed in the data: More precisely, if following replication both daughter cells contribute, on average, with the same statistical weight to the sequenced cell population, the net effect at the population level would be independent of the local DNA methylation level in the parent population. Since errors predominantly lead to a loss of DNA methylation at a probability proportional to the chance that a site was methylated before replication, its impact on the copying of hemimethylated DNA would merely lead to a monotonic decrease of the DNA methylation drift with the DNA methylation level in the parent population (cf. Fig. 2b).

**Figure 2.**
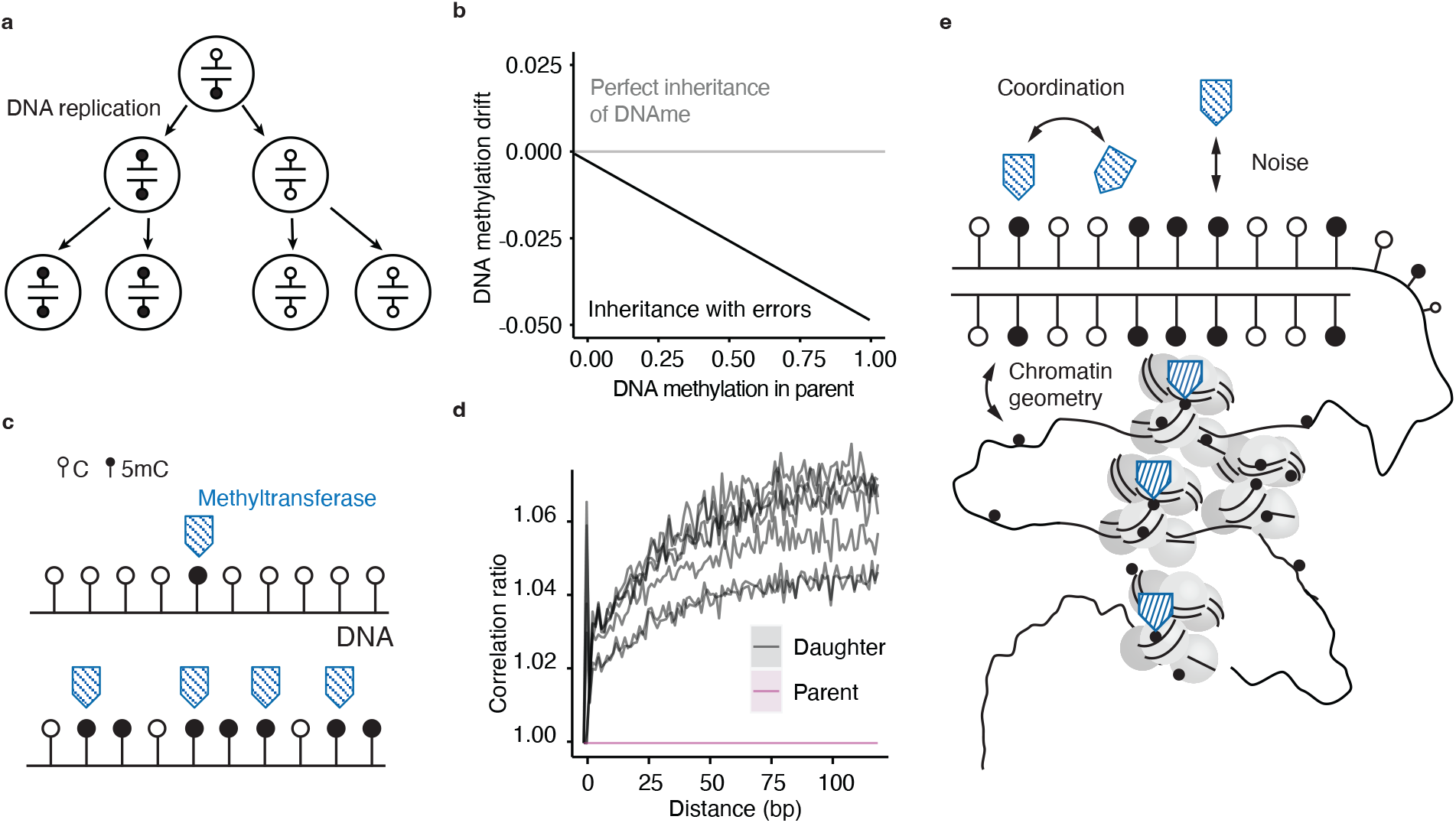
Minimal model for short-term DNAme drift. **a** Sketch of possible trajectories of hemimethylated sites after replication. **b** DNA methylation drift caused by the correction of hemimethylated sites during DNA replication. The negative slope of the DNA methylation drift is equal to the rate of errors in DNA maintenance per unit time. Higher error rates correspond to steeper lines as copy error-induced de-methylation happens faster. **c** Schematic of the effect of global variations in methyl transferase enzymes. A global decrease in methyl transferase enzymes (top) leads to a monotonic decrease in DNA methylation (everything else being equal) with the number of enzymes. Similarly, a global increase (bottom) would induce a global increase in DNA methylation. **d** Correlation of the DNA methylation state of pairs of CpGs as a function of the distance between these sites along the genome. The data shows the ratio between the correlation function of the daughters and the parent of the clonal MEF data (27). **e** Schematic illustrating the three components of the biophysical model. (For details, see main text.)

Therefore, in principle, the correction of hemimethylated sites could only explain the nonlinear drift observed in Fig. 1d under the condition of clonal competition between both daughter cells, such that a daughter cell with hemimethylated sites has a relative growth disadvantage and hence a lower statistical weight in the cell population. While this would lead to unstable fixed points of the DNA methylation drift on the population scale, to our knowledge, there is no reported evidence of a connection between the level of hemimethylation and the rate of cell proliferation.

Based on these considerations, we therefore reasoned that the observed nonlinear dependence of the DNA methylation dynamics likely originates from dynamics on the allele or cellular scale. On the cellular scale, changes in DNA methylation must ultimately be linked to changes in the global activities of methyl transferase enzymes (Fig. 2c). Changes in enzyme activity together with a non-monotonic dependence of catalytic rates on the DNA methylation level could potentially explain the DNA methylation drift in Fig. 1d. RNA-seq of the parent and daughter clones did not, however, show any difference in the expression of methyl transferase genes (27). Further, Chip-seq experiments show that the binding preference of DNA methylating and de-methylating enzymes depends monotonically on the DNA methylation level (31; 32).

Therefore, given the arguments above, we reasoned that the observed DNA methylation drift (Fig. 1d) must be a result of the accumulation of events on the molecular scale. In this case, feedback must happen between genomic positions on the same allele. To support the idea that DNA methylation changes are coordinated in space, we calculated the changes in spatial correlations between the DNA methylation values of pairs of nearby CpG sites. The degree to which genomic sites at a genomic distance *d* are correlated is quantified by the connected correlation *C*(*d*) between methylation states of all pairs of sites with distance *d*. To quantify correlative changes beyond uncorrelated noise, we computed the ratio of correlation values in daughters and parent cell cultures, *C*_daughter_(*d*)*/C*_parent_(*d*) (33). Our analysis showed that the change in DNA methylation between parents and daughters is positively correlated over distances of hundreds of base pairs (Fig. 2d). This suggests that CpG sites interact in such a way that nearby sites converge towards a shared DNA methylation level.

To test this hypothesis, we defined a minimal molecular-level model for the stochastic kinetics of enzyme-driven changes in the DNA methylation state of individual genomic sites. This model incorporates three basic components (Fig. 2e): first, spatial feedback of enzyme binding along the DNA sequence (indeed, several methyltransferases, including DNMT1 and DNMT3, have been shown to act cooperatively along the genome (34)); second, stochasticity, which arises naturally due to the random nature of enzyme binding and unbinding kinetics; and, third, the geometric organization of chromatin (as defined below). Transitions between different DNA methylation states depend on the DNA methylation levels of all other sites in proximity. Indeed, as shown by the strong and positive correlation in the dynamics between nearby sites (35) (Fig. 2d), this feedback is positive, leading to the tendency of nearby CpG sites to converge to similar DNA methylation levels.

Therefore, within the framework of this minimal model, the expected DNA methylation level, 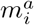, at a genomic position *i* on allele *a* evolves stochastically according to the equation,

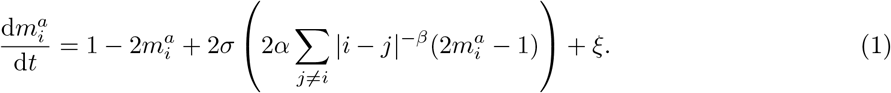

This Langevin-like equation incorporates at a “mesoscopic” level the three building blocks: The first term on the right-hand side of the equation describes the deterministic evolution of DNA methylation levels towards a common value of *m* = 1*/*2 due to the accumulation of stochastic local switching events. The second term encodes the positive feedback of DNA methylation levels between neighboring CpG sites and depends on a function *σ* with an argument that depends on a dimensionless coordination strength *α* and the integrated levels of DNA methylation weighted by proximity. Here, *σ*(*x*) is a function with sigmoidal form, reflecting the effect of the saturation of binding events when most neighboring sites are occupied (36) (Methods). Here, for simplicity, we set *σ*(*x*) = 1*/* (1 + *e*^*x*^). The factor |*i* − *j*|−*β* reflects the probability that two CpG sites with a genomic separation of |*i* − *j*| are in physical contact. The exponent *β* depends on the degree of chromatin compaction and takes typical values between 1 (for a fractal globule) and 2.1 (for a self-avoiding random walk) (37). Finally, *ξ* represents a noise term that takes random values at each instant of time and describes the fluctuations induced by random changes in DNA methylation states (Supplemental Theory). The Langevin-like dynamics of DNA methylation levels mirrors the dynamics of local magnetic moments in a classical ferromagnet. In this setting, the parameter *α* = *J/k*_*B*_*T* represents the ratio of a magnetic exchange energy, *J*, and the typical strength of thermal fluctuations, *k*_*b*_*T*. Here, the parameter *α* encodes the sensitivity of DNA methylation changes with respect to the DNA methylation state of the CpGs in the spatial proximity. It therefore quantifies the relative importance of spatial coordination and stochasticity in the DNA methylation kinetics.

### Test of model predictions

To investigate the validity of the model for the short-term DNA methylation drift, we tested its predictions against the results of sequencing experiments. The model predicts that the DNA methylation drift changes qualitatively as a function of the coordination strength *α*: if the coordination strength is weak compared to the magnitude of noise (*α <* 1), stochasticity dominates and the model predicts convergence to a configuration that maximizes the entropy leading to convergence towards a single stable fixed point at *m*^∗^ = 1*/*2. If the coordination strength is large, on the allele level, DNA methylation values tend to take similar values, leading to the emergence of two stable fixed points at *m*^∗^ = 0 and *m*^∗^ = 1 separated by an unstable fixed point at *m*^∗^ = 1*/*2 (Fig. 3a-c). In the language of dynamical systems, this abrupt change of the fixed point structure is called a bifurcation and, in the current context, a supercritical pitchfork bifurcation.

**Figure 3.**
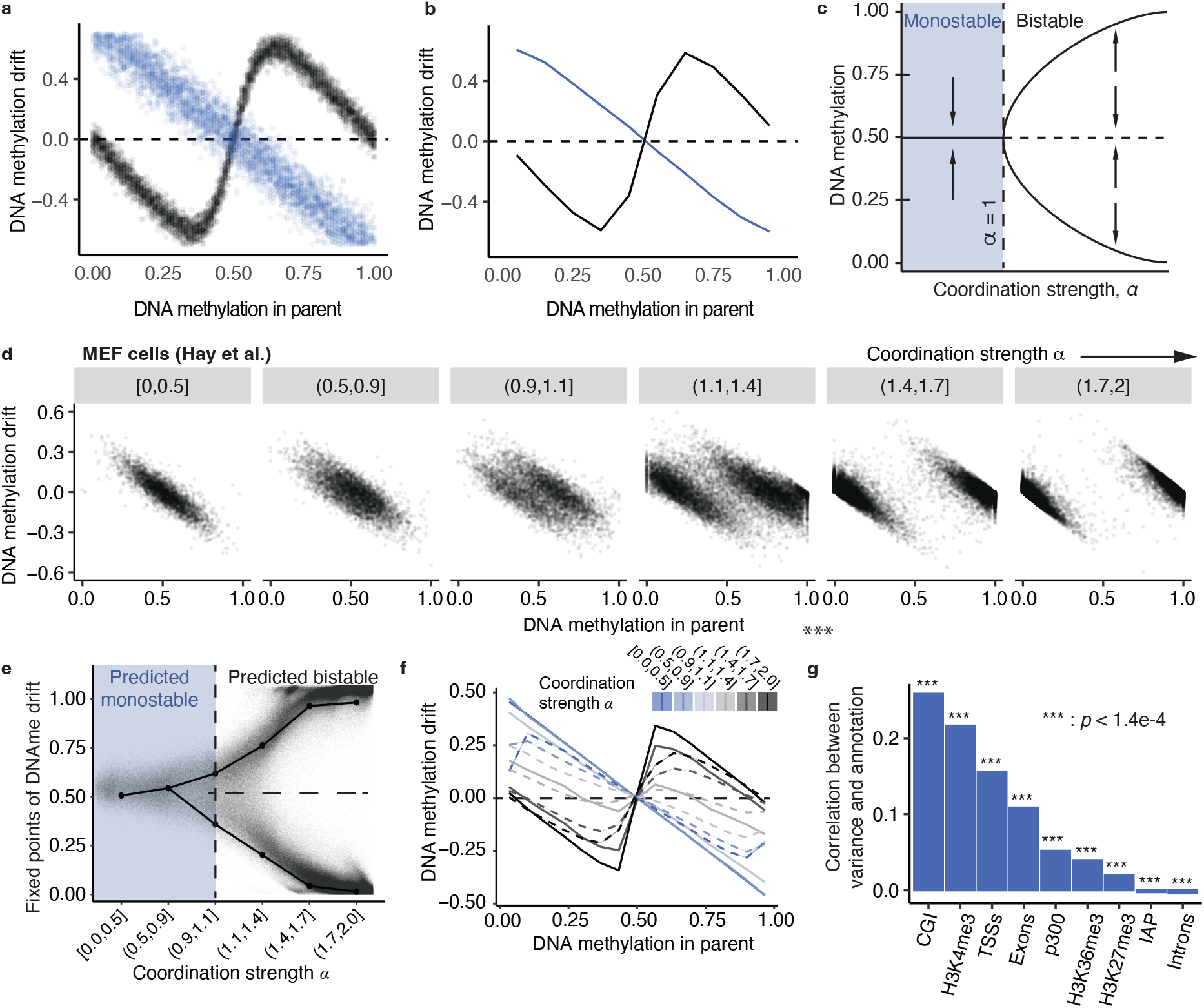
Comparison of theoretical and empirical DNA methylation drift. **a** Predicted DNA methylation drift for individual CpGs obtained from stochastic simulations of the feedback model of DNA methylation dynamics (1). Each dot represents one site. Color denotes sites with high values of the coordination strength (black, *α*=10) and low values of the coordination strength (blue, *α* = 0.5). **b** Average DNA methylation drift obtained from the same stochastic simulations. Parent DNA methylation levels were grouped into 10 bins. **c** Predicted dependence of the number and stability of fixed-points depending on the coordination strength *α*. Solid horizontal lines represent stable fixed points, and dashed horizontal lines represent unstable fixed points. The vertical dashed line marks the bifurcation points. **d** Empirical DNA methylation drift for different values of the coordination strength in the clonal MEF dataset (27). Each point corresponds to a covered CpG site. **e** Position of stable fixed-points inferred from experimental data (points and solid lines, see Methods) and daughter DNA methylation levels as a function of the inferred coordination strength. Each dot represents a covered CpG site. **f** Average DNA methylation drift for sites grouped by different values of the inferred coordination strength *α*. Solid lines indicate the predicted DNA methylation drift from stochastic simulations, dashed lines represent experimental data. **g** Pearson correlation between the inferred values of *α* and several genomic annotations for the clonal MEF data (27).

To compare the model predictions to the sequencing data, we estimated the parameter *α* at each genomic locus by using a simulation-based inference method (38) (Methods). Specifically, parameter estimates were determined using aggregate statistics from stochastic simulations of the model (1) to infer the most probable value of *α* that yielded a given mean and variance of the daughter DNA methylation for a given value in the parent. Based on this, we then estimated the value of *α* for each covered CpG site. By computing the empirical DNA methylation drift separately for every site with different inferred values of the coordination strength *α*, we found that the DNA methylation drift indeed exhibits the same fixed point structure as a function of *α* as that predicted by the model (Fig. 3d). Moreover, further analysis of the data showed evidence of the predicted supercritical pitchfork bifurcation with increasing values of *α* (Fig. 3e). Finally, when initialized with the site-specific DNA methylation values in the parent populations, the stochastic simulations of the model also quantitatively predicted the profiles of DNA methylation drift for given values of the coordination strength *α* (Fig. 3f).

The model describes the genomic feedback between nearby CpG sites with a strength that depends on the coordination parameter *α*. It, therefore, follows that genomic regions with high inferred values of *α* are more spatially correlated than regions with low inferred values of *α*. To test this prediction, we calculated the spatial correlation function, which encodes the tendency of a pair of sites to have the same DNA methylation level as a function of the distance between these sites. Consistent with the prediction of the model, we found that sites with high values of the inferred coordination strength *α* were more strongly correlated than sites with low values of *α* (Fig. S1a). As the majority of sites are assigned *α >* 1 (Fig. S1b), this indicates a tendency of DNA methylation to align with the neighboring methylation context. Moreover, the dynamics obtained from the simulation of the model reproduced the non-linearity observed in the dependence of DNA methylation drift on parent DNA methylation ((Fig. S1c)) and the transition in daughter DNA methylation distribution at different values of *α* (Fig. S1d,e). Together, these results further support our hypothesis that DNA methylation dynamics involves spatial coordination across proximate CpG sites.

The qualitative change in DNA methylation dynamics as a function of the coordination strength *α* and its inference from the analysis of data from the short-term sequencing study allows us to rigorously classify CpG sites into those with stable epigenetic memory (*α <* 1) and those that follow a more stochastic dynamics (*α >* 1). To quantify the relative distribution of more “deterministic” and “stochastic” sites, we quantified the association of *α* with different genomic annotations for MEFs (Fig. 3g). The results showed that high values of *α* were particularly associated with regions involved in epigenetic and genetic regulation, such as CpG islands (CGIs), transcription start sites (TSSs), and expression regulation involving histone modifications such as H3K4me3. In contrast, H3K27me3, IAP, and intronic regions were only weakly correlated with *α*. This suggests a high control of DNA methylation dynamics for genomic regions associated with active chromatin.

By focusing on experimental data covering DNA methylation dynamics on the timescale of a few rounds of cell division, these results show how the kinetics of enzyme binding to the DNA lead to a convergence of DNA methylation concentrations towards steady-state levels. This short-term dynamics of DNA methylation levels is captured quantitatively by a minimal phenomenological model, characterized by its fixed-point structure and a supercritical pitchfork bifurcation that takes place as a function of a single dimensionless parameter, *α*, quantifying the local strength of spatial coordination.

### From short-time DNA methylation drift to aging dynamics

From a dynamical systems perspective, the number of fixed points, their stability, and their transition through the supercritical pitchfork bifurcation uniquely characterize the dynamics of DNA methylation on short timescales. By comparing the fixed-point structure and their bifurcations between the short-term experiments and the long-term aging dynamics, the feedback model allows us to test whether the same dynamical system drives both processes. To this end, we sought to determine these features using reduced representation bisulfite sequencing experiments on blood, cortex, heart, lung, and liver tissue, which provide high coverage of DNA methylation at the population level (39; 28). We complemented these bulk experiments with single-cell bisulfite sequencing experiments on mouse blood (29), which provide lower coverage but allow for an estimate of the relative coordination strength, *α*, at the single-cell level.

In the context of aging, we defined DNA methylation drift as the difference in DNA methylation levels between consecutive time points. To estimate the parameter *α* in the aging datasets, we again used a simulation-based inference method (Methods). As before, we inferred the site-specific value of the coordination strength *α* by comparing the experimental and theoretical means and non-technical contributions to the variances of DNA methylation levels. For the single-cell experiment, the variance in DNA methylation was calculated across single cells at a given time point, and we considered the dynamics of DNA methylation averaged over 100 bps (see Methods). For the bulk experiments, it was calculated across biological replicates. Based on these analyses, we then grouped sites by similar values of the coordination strength *α* and calculated the respective DNA methylation drifts. We found that, genome-wide, the DNA methylation drift during aging followed the same pattern as that observed on short timescales (Fig. 4a, b, Fig. S2a, b, c, d). Specifically, the DNA methylation dynamics during aging generally converged to the same fixed points as in the clonal MEF data (Figs. 3d-f) and exhibited the same supercritical pitchfork bifurcation structure with increasing values of *α*. Since the values of *α* for a given site did not change systematically during aging (Fig. S3a), we reasoned that the drift in DNA methylation is driven intrinsically by endogenous self-organization dynamics and not by extrinsic changes in the parameter defining the model. These results indicate that the dynamics of DNA methylation during aging are described by the same dynamical system as the fast dynamics that take place on timescales of a few cell divisions in the culture setting. It therefore follows that the dynamics of DNA methylation on the intermediate timescales of aging (the timescale between successive time points in the experiments) arises from the accumulation of individual molecular events that take place on the timescale of milliseconds.

**Figure 4.**
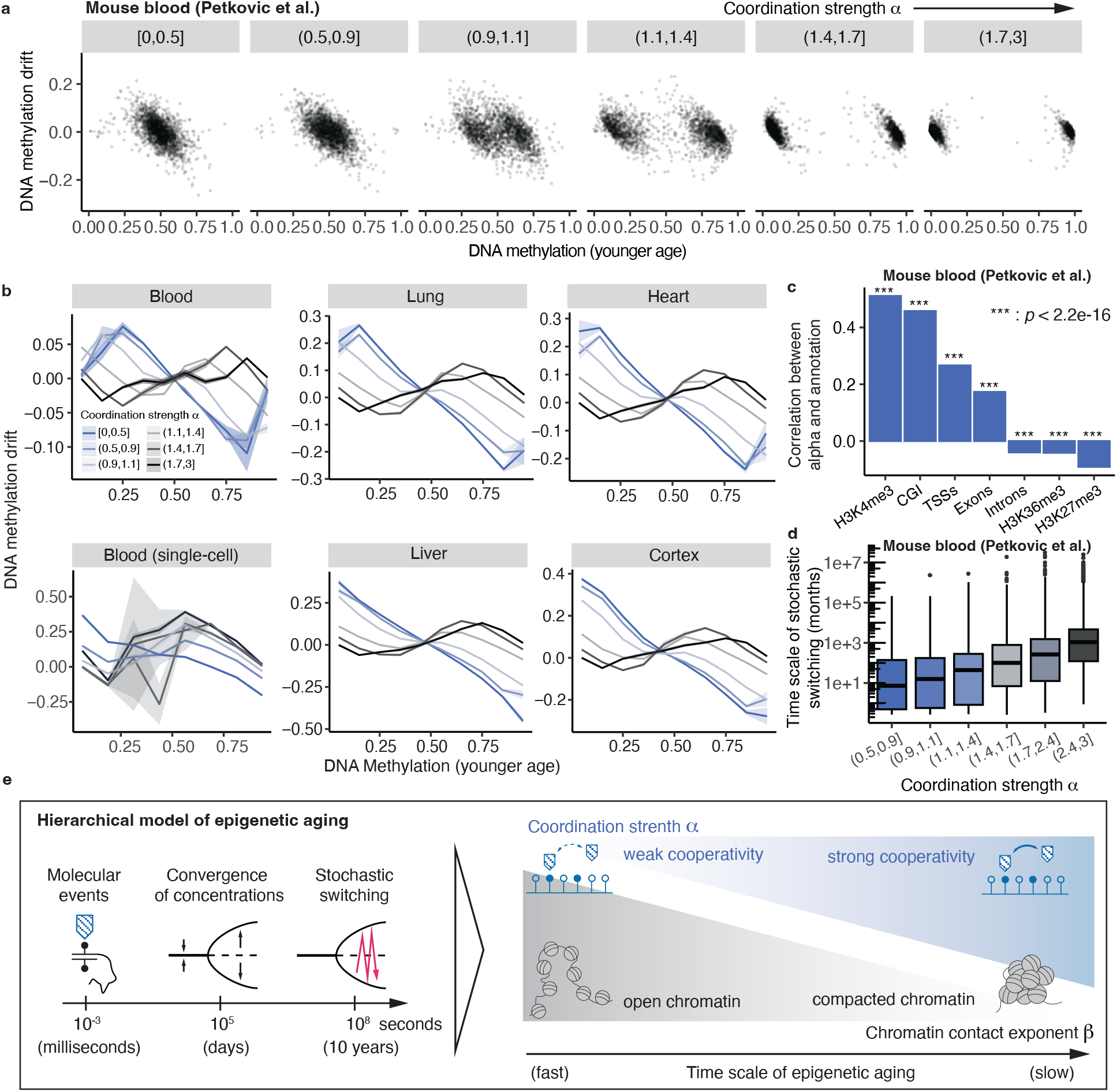
DNA methylation dynamics during aging. **a** DNA methylation drift for mouse blood (39) for different values of the inferred coordination strength *α*. Each dot represents a CpG site. **b** DNA methylation drift for different tissues from bulk sequencing (39; 28) and single-cell sequencing (29) data. For every experiment, the DNA methylation drift at a certain age is calculated as the difference of DNA methylation between the considered age and the previous one covered in the respective experiment (x-axis, 10 bins for bulk, 8 bins for single-cell) for different inferred values of *α*. **c** Spearman correlation between the inferred values of *α* and several genomic annotations for mouse blood from animals of age 24 months (39). **d** Box plot of the inferred aging time scale for different ranges of the inferred coordination strength *α*. Since there is no discernible drift towards 1*/*2 for sites whose DNA methylation level was on average in the interval [0.4, 0.6], we removed these sites from the analysis. Horizontal lines represent the median, the lower and upper hinges correspond to the first and third quantiles, and whiskers represent 1.5 times the interquartile range. The inset shows the Spearman correlation coefficient and corresponding p-value for the Spearman correlation between the Age-average *α* value and the inferred aging scale. **e** Graphical summary of the hierarchical model of DNA methylation aging. DNA methylation dynamics is characterized by a hierarchy of time scales. This hierarchy is determined by an interplay of coordination strength between methylation events and chromatin structure.

We next questioned whether, during aging, the genomic association of sites with high or low values of coordination strength is consistent with the results obtained from the study of the short timescale dynamics (Fig. 3g). To this end, we again correlated the overlap of sites with genomic intervals given by different genomic annotations with the value of *α* estimated for a given site (Fig. 4c). We found that the values of *α* were largely consistent in that active genomic regions, such as H3K4me3-annotated regions, are associated with a high coordination strength, epigenetic memory, and a more deterministic time-evolution. This further supports the conclusion that DNA methylation dynamics on short and long timescales are driven by the same mechanism.

From these results, we conclude that the dynamics of DNA methylation on intermediate timescales result from the cumulative impact of fast molecular processes. The timescale on which these changes occur during aging is governed by the convergence time of DNA methylation levels to the fixed points of Eq. (1), i.e., scaling in proportion to the inverse of the coordination strength 1*/α*. The timescale associated with DNA methylation aging is also dependent on the local chromatin structure through the exponent *β*, characterizing the internal end-to-end distance distribution of genomic loci (37). Small values of *β* (viz. closed chromatin) show a faster convergence to the fixed point than larger values of *β* (open chromatin).

### Quantification of the slowest time scale of aging

On timescales much larger than the time it takes for DNA methylation levels to converge to a steady state, noise may lead to stochastic transitions between the stable fixed points of Eq. (1) and an ensuing overall convergence of DNA methylation levels to a value of 1*/*2, a result confirmed by stochastic simulations (Fig. S3b, Methods). Indeed, as has been observed previously (39), we found evidence of a general convergence of DNA methylation levels towards 1*/*2 (Fig. S3c, Methods). Moreover, as predicted by the model, the rate of convergence to a value of 1*/*2 is faster if the effects of stochasticity are strong compared to genomic coordination, scaling in proportion to the inverse value of *α* (Methods). To estimate the timescale on which such stochastic switching becomes dominant, we first derived the effective drift of the average of DNA methylation levels across replicates (Methods), 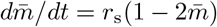, where *r*_s_ is the rate of switching between the fixed points (40). By solving this equation, we used linear regression to estimate the timescale 1*/r*_s_ associated with this process. We estimated that, over a timescale of roughly 1000 months (80 years), stochastic switching between stable fixed points dominates DNA methylation dynamics (Fig. 4d). This is much longer than the lifespan of mice, but comparable to the lifespan of humans.

Altogether, these results suggest a hierarchical model of aging in which DNA methylation aging does not emerge from a slow process driven by external factors, but from the self-organized accumulation of fast molecular events (Fig. 4e). This accumulation manifests in different ways across temporal scales: On intermediate timescales (days to weeks), DNA methylation aging is dominated by the convergence of local DNA methylation levels to fixed points given by a mechanism that combines local coordination of genomic loci and noise. On much longer time scales (80 years), stochastic switching between these fixed points dominates and leads to a slow convergence towards DNA methylation levels of around 1*/*2.

## Discussion

Previous studies have emphasized how changes in DNA methylation can encode information on aging. However, the link between slow, deterministic changes that occur in DNA methylation over the long term and the stochastic molecular events that take place over much shorter timescales, as well as the molecular mechanisms that drive these associations, has remained in question. Since DNA methylation can be profiled with single-base pair resolution and changes systematically during aging, it provides a suitable context to address this question. To bridge temporal scales, we took advantage of two experimental data sets that each tackle a different timescale of DNA methylation dynamics. The clonal MEF data provided a bridge between the timescale of individual molecular events (milliseconds) and the timescale of cell divisions (days in MEFs) on which DNA methylation concentrations converge towards their steady-state levels. Based on these findings, we showed that the short-term dynamics of DNA methylation could be largely captured within the framework of a minimal model, based on a single adjustable parameter. Finally, using longitudinal data during aging, we showed that the same model could capture the long-term dynamics of DNA methylation.

From these results, we conclude that there is a three-fold hierarchy of processes that dominate DNA methylation aging that occur on different timescales: First, there are individual stochastic events in which enzymes interact with the DNA and each other. These dynamics are characterized by molecular binding rates and occur on the timescale of milliseconds. Second, there is a convergence of DNA methylation concentrations towards the steady state. The rate at which aging occurs on these scales is defined by the strength of genomic coordination of DNA methylation and the local chromatin architecture (days to months). Third, the accumulation of rare events leads to stochastic transitions between these steady states. The rate of these transitions depends exponentially on the inverse of the coordination strength and the typical genomic size over which DNA methylation is correlated. It therefore gives rise to slow DNA methylation dynamics over the timescale of decades.

Here, to gain a conceptual understanding of how temporal scales of DNA methylation are bridged, we placed emphasis on a minimal phenomenological model of DNA methylation dynamics, relying on the arrangement of fixed points within dynamical systems theory. Nevertheless, there are plausible molecular mechanisms for the different processes that comprise the model. The DNA methyl transferases DNMT1 and DNMT3 have been shown to act cooperatively and processively over genomic domains of 100bps or 500bps, respectively (30; 41). Moreover, the emergent length scale of the change in correlations between genomic sites in Fig. 2e suggests a DNMT1-related mechanism.

Taken together, our work suggests a hierarchical model of aging in which molecular self-organization processes give rise to different dynamics on distinct spatial scales.

## Acknowledgements

We thank Anne-Ferguson Smith and Amir Hay for sharing their experimental data prior to its publication as well as illuminating and encouraging discussions at an early stage of our work. B.D.S. acknowledges the support of the Wellcome (220379/B/20/Z) and the Royal Society through an EP Abraham Research Professorship (RSRP\R\231004). This project has received funding from the European Research Council (ERC) under the European Union’s Horizon 2020 research and innovation program (grant agreement no. 950349).

## Author Contributions

M.C., B.D.S. and S.R. conceptualized the project. M.C. performed data analysis, theory, and simulations. M.C., B.D.S., and S.R. wrote the manuscript.

## Data Deposition Statement

Published data sets used in the analysis are listed in Supplementary Table 1.

## Methods

### Derivation of the model describing DNA methylation drift

In this section, we derive the kinetic equation (1) for the average DNA methylation at a given genomic position. We derive this equation by starting from a stochastic model incorporating individual molecular binding events. We then derive a coarse-grained stochastic description by performing an expansion in the system size. To begin, we consider a segment of the DNA, which is represented by a one-dimensional lattice of size *N*. The DNA methylation state on each site *i* is represented by a binary random variable *s*_*i*_ ∈ {−1, 1} with *i* ∈ {1, *N*}. These variables represent methylated (*s*_*i*_ = 1) and unmethylated (*s*_*i*_ = −1) CpG states.

We describe the time evolution of the DNA methylation states as a Markov process. A Markov process is fully defined by transition rates, which give the conditional probabilities of the system transitioning from one state to another per unit time. Our data and previous publications suggest that DNA methylation events are locally correlated, such that the transition rates depend on DNA methylation values in the vicinity of a given site *i*. This dependence is mediated by the local conformation of chromatin. This is statistically described by the inner contact probability, which decays algebraically with an exponent *β*. Specifically, the contact probability takes the form *C Σ*_*j≠i*_ |*i* − *j*|^−*β*^, where *C* is a normalization factor that ensures that the sum is normalized to 1.

Given that two sites *i* and *j* are in contact, the tendency of a transition to occur is proportional to *s*_*i*_*s*_*j*_, such that if sites *i* and *j* have an unequal DNA methylation state, transitions to the opposite state are promoted. With this, the transition rate *W* (−*s*_*i*_|*s*_*i*_) at which a state *s*_*i*_ at site *i* transitions into its opposite state −*s*_*i*_ (i.e., methylated to unmethylated or *vice versa*) reads

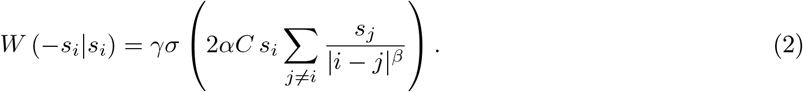

Here, *α* is a dimensional parameter governing the strength of coordination. In thermodynamic equilibrium, *α* is the ratio of the interaction energy and the energy of thermal fluctuations, *k*_*B*_*T*. The parameter *γ* is a rate that denotes the number of state transitions per unit time. *σ*(*x*) describes the effect of saturation when a large fraction of sites in a given DNA segment have identical DNA methylation values. It is a sigmoidal function varying smoothly between 0 and 1. Here, we chose the sigmoidal function 1*/*(1 + *e*^*x*^) as in Ref. (36). This definition of the transition leads to dynamics in which sites will preferentially transition to values aligning with the most common DNA methylation state in the system. The parameter *α* sets the steepness of the sigmoidal function and, consequently, the sensitivity of transition rates to changes in the average methylation. With these transition rates, the time evolution of the probability of finding the system in a given configuration {*s*_*i*_} follows a master equation of the form

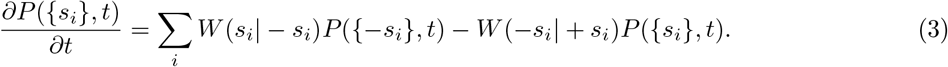

The exponent *β* takes values between 1 for space-filling chromatin and 2.1 for self-avoiding random walks. For exponents *β <* 3*/*2, the mean-field theory of the Markov process defined by the transitions is valid. This compares with typical values of *β* in cells determined from Hi-C experiments, which predict values around unity. We therefore apply the mean-field limit when considering the spatial dependence of the interactions. In this case, the transition rates read

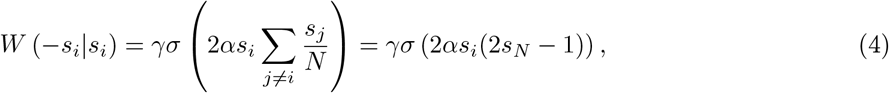

where we have introduced the average site occupation variable *s*_*N*_ = *Σi*(1 + *s*_*i*_)*/*2*N* that counts the number of methylated sites in the DNA segment. For the normalization chosen, different values of *β* will have the same critical *α* value.

In this limit and for large system sizes, *N* ≫ 1, we can substitute the spatial average *s*_*N*_ with the ensemble average 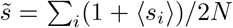, where 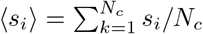 denotes the single-site ensemble average and *N*_*c*_ is the number of cells included in the experiment. From these approximations, we obtain the following expression for the time-evolution of the probability of finding a given ensemble average 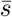 at a given genomic position,

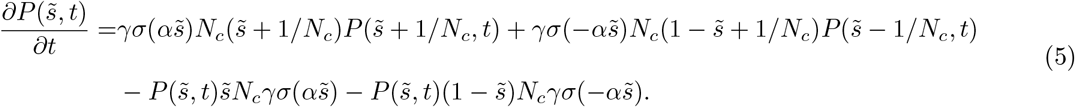

The first line of the right-hand side of Eq. (5) represents the contribution of transitions to the average 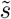, while the second line corresponds to transitions either decreasing or increasing the average methylation 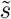. By expanding this equation in 1*/N*_*c*_ and retaining terms of order O(1) for the drift and O(1*/N*_*c*_) for the diffusion term, we obtain the Fokker-Planck equation,

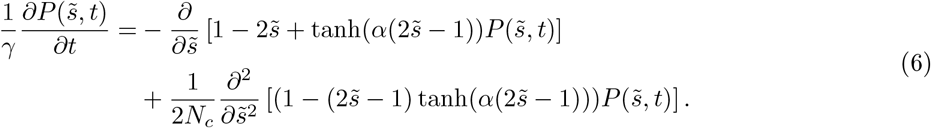

This equation is equivalent to a Langevin equation for the average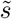,

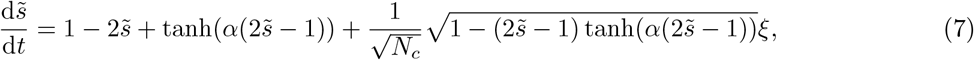

where *ξ* is a Gaussian white noise with correlation ⟨*ξ*(*t*)*ξ*(*t*^′^)⟩ = *δ*(*t* − *t*^′^) and the equation is interpreted in the Itô sense (42).

In this framework, the deterministic part of the time evolution of the average DNA methylation in a genomic region across cells follows from Eq. (7) by setting the noise to zero,

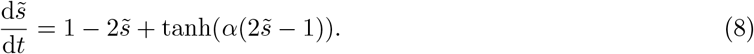

In this model, the time-evolution of DNA methylation values is therefore determined by a single parameter, *α*, which controls the balance between stochasticity and genomic coordination. High values of *α* characterize genomic sites where DNA methylation is maintained with high fidelity, providing epigenetic memory, while low values of *α* characterize sites where DNA methylation changes are dominated by noise. Mathematically, the right-hand side of Eq. (7) gives the stochastic DNA methylation drift and it exhibits two stable fixed points given by its roots at 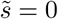 and 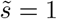, and an unstable fixed point at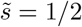.

### Stochastic simulations and estimation of the coordination strength

Equation (7) was simulated through a Murayama-Euler first-order scheme for stochastic differential equations. We set *N*_*c*_ = 32, translating to the median coverage per site over the MEF dataset. To estimate the value of the coordination parameter *α* for a given CpG site, we used the simulation-based inference toolkit in Python (38). This method uses aggregate statistics from stochastic trajectories to infer the best set of parameters producing the observed dynamics. For the parameter inference, Eq. (7) was simulated for 10^3^ time steps with the time increment set to *dt* = 0.01, one for each allele for a total of two simulated trajectories. From these results, we calculated the corresponding average and variance of methylation values at the final time point. We then trained a neural network with the SBI-toolkit to infer the joint posterior distribution of *α* values and initial average DNA methylation values for a given combination of the mean and variance of DNA methylation values at the end of the simulation, starting from a prior of parameters for which *α* ∈ [0.0, 2.0] (38). From these results, we then took the MEF data and a sampled value of *α* from the posterior distribution obtained above for every CpG site and cell line for the given empirical values of the mean and variance of DNA methylation values over daughters. To compare the predicted DNA methylation drift of the model to the experimental data in Fig. 4b, we then simulated seven trajectories for each covered CpG site with the estimated value of *α* and with initial conditions taken from the parent DNA methylation in the MEF experiment.

The theoretical predictions of the DNA methylation drift in Figs. 3a-c were obtained by drift estimates from random initial conditions with Gaussian white noise with variance 0.01 for high coordination strength and 0.2 for low coordination strength. Line plots in Fig. 3b are obtained from the scatter plots in Fig. 3a by computing the binned average over the initial DNA methylation values.

To estimate the values of *α* in the aging datasets, we again ran an inference procedure of the parameter *α* with the sbi toolkit (38). The estimation for the values of *α* was limited to the interval [0.0, 3.0] and, in addition to the average and variance of DNA methylation (as considered in the MEF data), we considered as input parameters of the model the single-site coverage *N*_*c*_ as well as a proxy of the number of cells whose dynamics is probed, with prior *N*_*c*_ ∈ [0, 1000]. This allowed us to use the coverage as input for the model dynamics while analyzing a wide range of coverage values. The posterior was inferred from the aggregate statistics of the average and variance of DNA methylation over five simulated single-cell trajectories, corresponding to the average number of replicates in the different datasets. For the inference of *α* on the aging data, we used the average variance of DNA methylation and the average coverage over biological replicates for every site and age as input for the posterior sampling. We inferred the value of *α* for 100,000 randomly sampled sites in the Stubbs and Petkovich datasets (28; 39) for sites with coverage lower than 1000 (to stay within the bounds employed for the prior distribution). For the single-cell blood dataset, given the low coverage at every site, we inferred the value of *α* by simulating the dynamics of the DNA methylation average over windows of 100bps. For the inference procedure, we considered *N*_*c*_ = 100, as we now consider the average over 100 CpG sites.

### Transition rates between fixed points of the average DNA methylation dynamics

In this section, we derive an expression for the transition rates between the attractors of the DNA methylation dynamics, as given by the stochastic differential equation (1). Here, we refer to (40) for the derivation of the transition rates between attractors in the presence of multiplicative noise. In our case, the transition rate reads

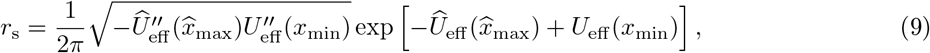

with effective potentials given by

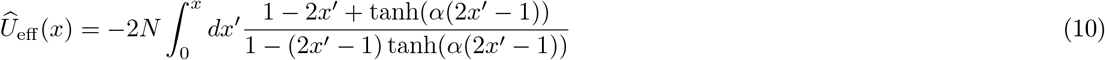

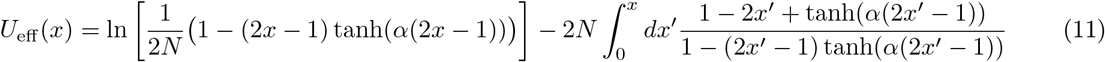

and *x*_min_, 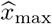 correspond respectively to the minimum of *U*_eff_^′^ and the maximum of *Û*_eff_.

In contrast to the rate decay to the fixed points, transitions between fixed points are suppressed by the exponential of the difference between the maximum and the minimum of the two effective potentials defined above. These transitions consequently happen at a lower rate compared to the convergence to fixed points. In particular, the rate of transition is suppressed exponentially by the number of interacting sites *N* and the barrier height between them *U*_eff_ and *Û*_eff_. Note that this barrier decreases with decreasing values of *α*, such that the typical time for transitions is proportional to the coordination strength.

The rate of change of the DNA methylation average close to the fixed point (here we consider the fixed point at *m* = 0 as an example) is given by the linearized equation of motion for the average DNA methylation value

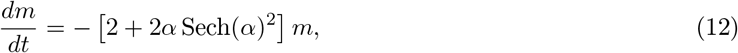

whose solution *m*(*t*) = *m*(0) exp −*t*(2 + 2*α* Sech(*α*)^2^)] shows an exponentially fast relaxation towards the fixed point.

### Estimation of the slowest timescale of aging from linear regression of DNA methylation dynamics

In this section, we derive the equation for the average DNA methylation on aging timescales. To this end, we note that on timescales larger than the ones associated with convergence to the fixed points of Eq. (8), the dynamics is dominated by noise-induced transitions between the different attractors. These transitions happen at a certain rate *r*_*s*_ that depends on the strength of the fluctuations and the barrier between the attractors (see previous Methods section). On average, the probability *p*_1_ of observing the state around the attractor at high DNA methylation then follows the equation,

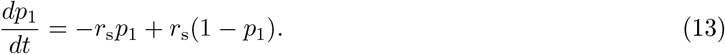

The first term on the right-hand side describes the probability outflux due to transitions from methylated to unmethylated states, while the second term describes the probability influx from unmethylated states. As the probability of being in the methylated states corresponds to the average DNA methylation, Eq. (13) can be integrated to give the drift of the population average DNA methylation as

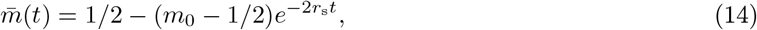

where *m*_0_ is the initial population average. At the onset of the aging drift dynamics (i.e., for small times *t*), this expression can be expanded as 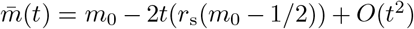. From this result, one can obtain the expression for the timescales over which aging becomes relevant from a linear regression on the DNA average dynamics 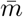, where the regression slope *b* is related to the aging timescales through the relation 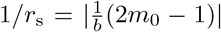. For the linear regression in the Petkovich dataset (39), we used this method on the randomly sampled 100,000 sites by fitting the linear regression on the methylation time-dependence from age 12 months, without considering sites for which the DNA methylation at that age was not sampled and sites for which 11 ages were retained.

### Analysis of sequencing data

For MEF data from (27), DNA methylation drift was calculated as the difference between the daughters’ average DNA methylation and the parent’s DNA methylation for every CpG site. Daughter variance was calculated as the variance of methylation levels among daughters.

Spatial correlation functions were calculated as the connected correlation coefficients by calculating the connected correlation between CpGs at fixed distances up to 3000 bps. Fixed points of the dynamics of DNA methylation levels were obtained as modes of the average daughters’ DNA methylation distribution for different levels of the coordination strength (Fig. 3e).

For the Stubbs and Petkovich datasets (28; 39), the average and variance of DNA methylation at single sites were obtained as the average and variance of DNA methylation levels between biological replicates for every site. DNA methylation drift was calculated as the difference in these average methylation levels between two successive time points at each site.

To compute the relative drift of DNA methylation for the Petkovich dataset (39), we focused on DNA methylation measurements taken after Age 12 months, as the DNA methylation drift distribution stabilizes from that age (Fig. S3(d)).

To compute the correlation between genomic annotations and the values of the coordination strength, we first assigned to every CpG site a binary variable with value of 1 or 0 depending on whether the site belonged to the chosen annotation or not. We then calculated for every type of region the Spearman correlation coefficients over CpG sites between the inferred *α* values and the annotation binary variable described above.

To check for possible dependence on coverage of the observed DNA methylation dynamics as a function of *α*, we analyzed the behavior of the DNA methylation drift as a function of *α* for different coverage thresholds. For the MEF dataset (27), the same behavior is observed for coverage thresholds of 30, 50, 100, and 200. For the Petkovich and Stubbs datasets (39; 28), the same behavior is observed for coverage thresholds of 30, 50, 100.

## Supplementary figures

**Figure S1.**
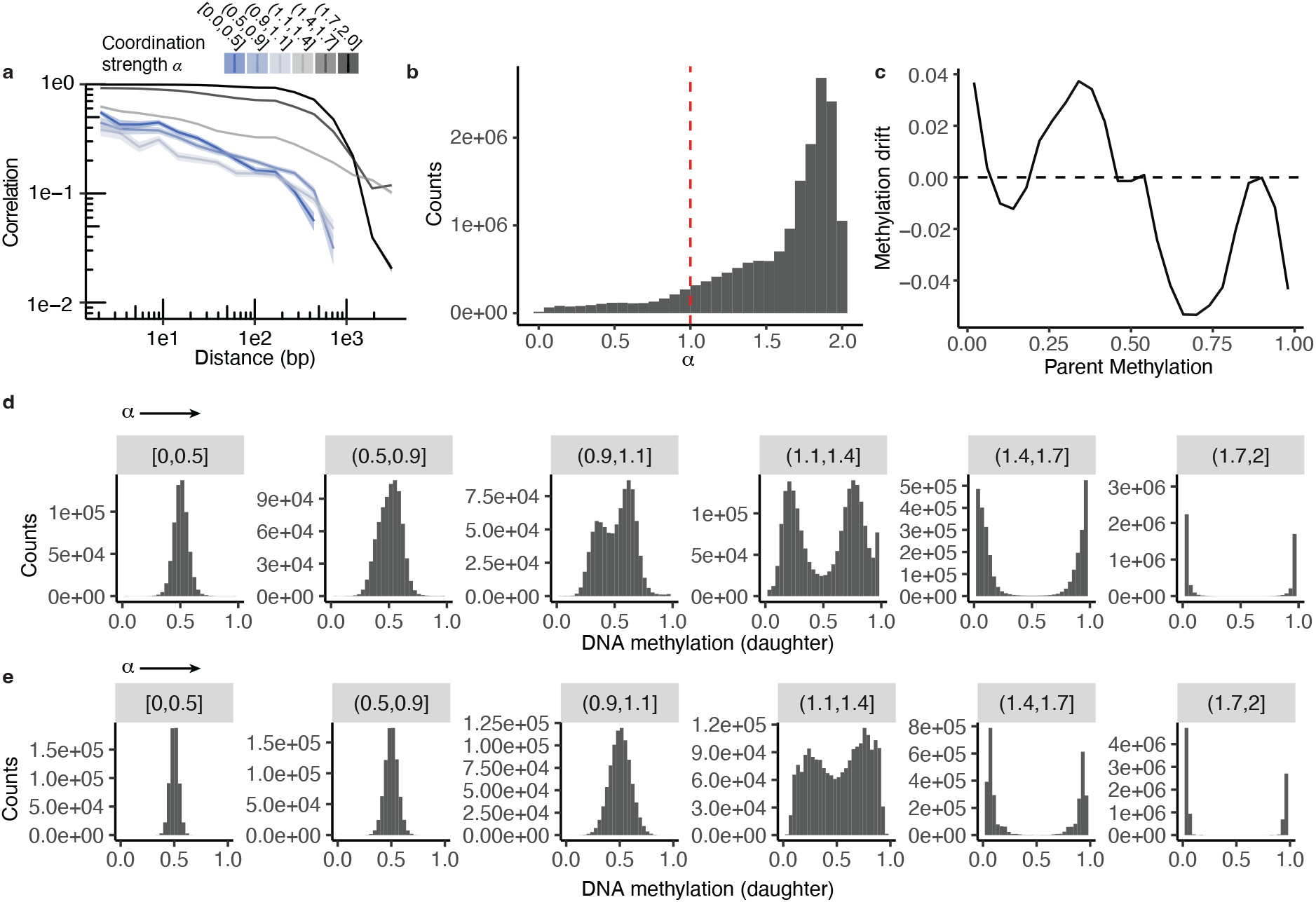
Estimation of the coordination strength for MEF cells. **a** Pearson correlation function of DNA methylation at different genomic distances and grouped by values of *α*. **b** Genome-wide histogram of inferred values of *α*. The red dashed line indicates the value of *α* corresponding to the bifurcation point. **c** Aggregate DNA methylation drift as a function of parent DNA methylation (25 bins) from stochastic simulations of the model values of *α* taken from the experiments. **d** Histograms of experimental daughter population average DNA methylation values (27) in daughter colonies for different values of *α*. **e** Histogram of daughter population average DNA methylation values as obtained from stochastic model simulations for the empirical values of *α*.

**Figure S2.**
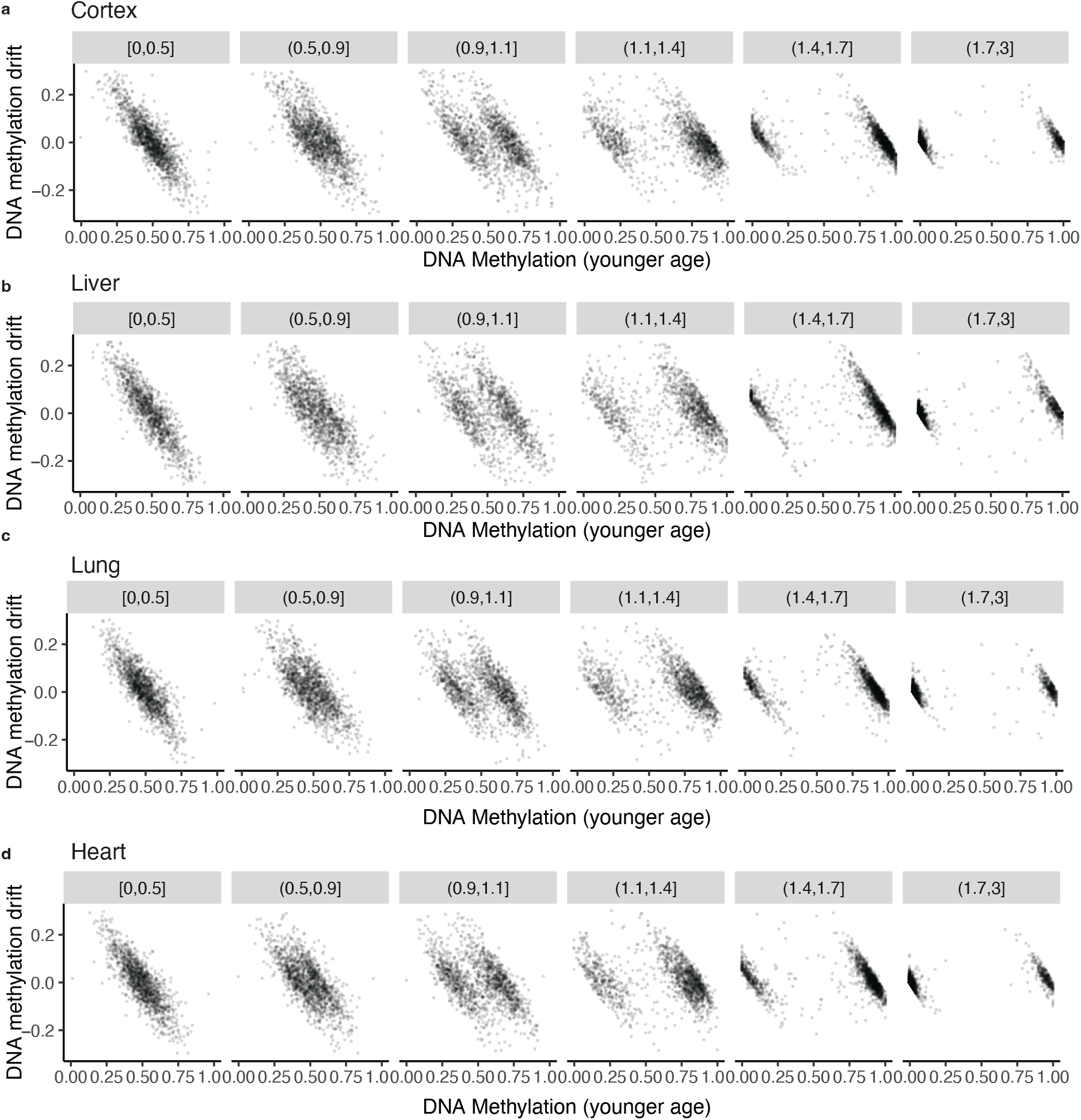
DNA methylation dynamics during aging. **a**-**d** DNA methylation drift for different intervals of the coordination strength *α* (columns) for different tissues in the Stubbs dataset (28). Points correspond to the CpG sites (100,000 sampled randomly)

**Figure S3.**
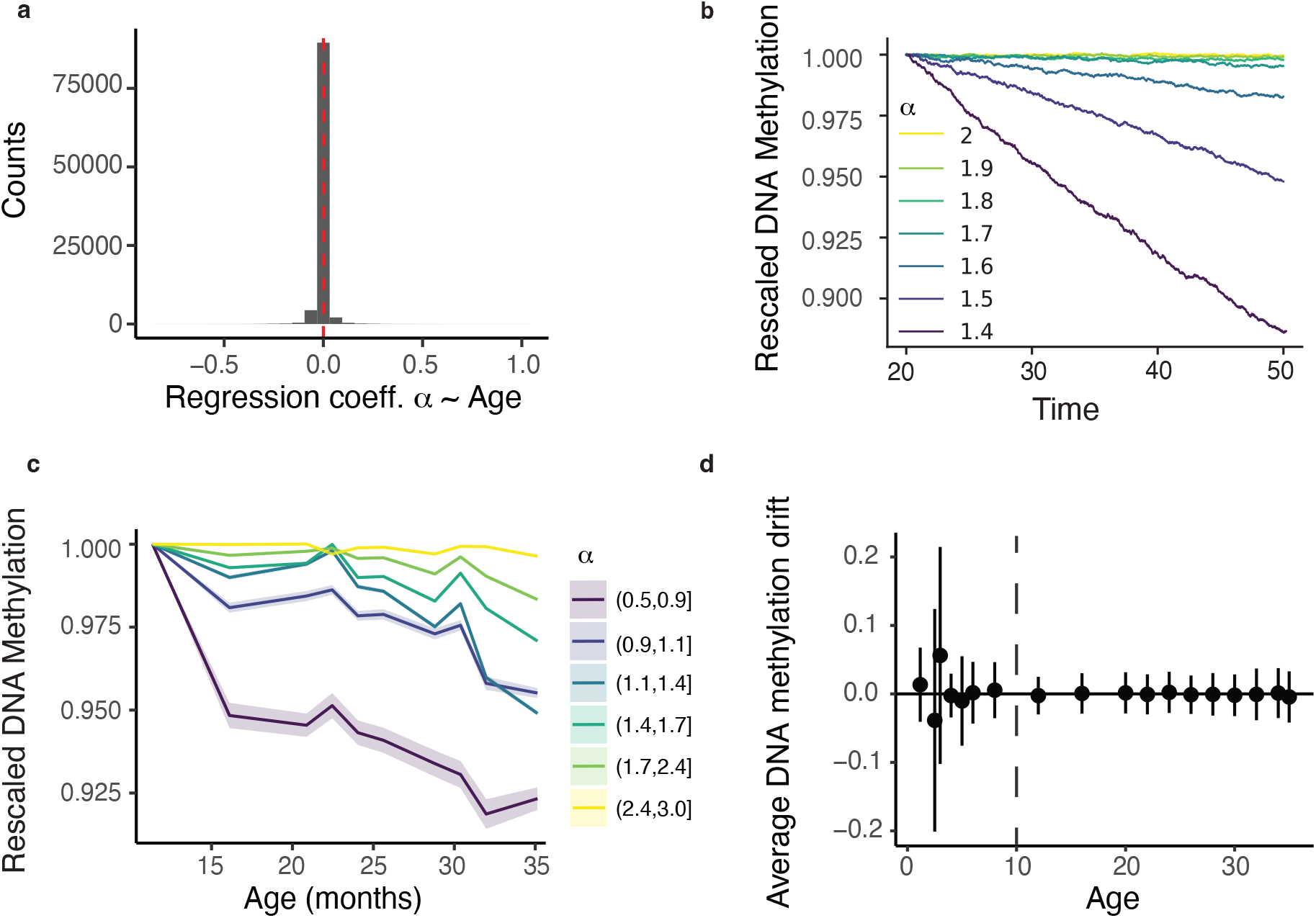
Drift of DNA methylation during aging. **a** Histogram of the regression coefficients obtained from a linear regression between the inferred values of *α* and age in the Petkovich dataset (39). The red dashed line corresponds to a regression coefficient of zero. **b** Dynamics of the simulated DNA methylation average over 1000 cells as a function of time and for different values of *α* (Methods). DNA methylation values were rescaled by the DNA methylation at the initial time point. **c** Time-evolution of the average DNA methylation of DNA methylation from Ref. (39) for different values of *α*. DNA methylation values in the interval *m* ∈ [0.0, 0.5] were transformed according to the formula *m* →1− *m*. In this way, low methylation values with increasing methylation show a decreasing trend similar to highly methylated sites. Furthermore, DNA methylation values were normalized by their value at 12 months. **d** Average DNA methylation drift as a function of age for Petkovich dataset as a function of age. The average is taken over CpG sites and replicates measured at a given age. Vertical bars indicate the standard deviation of the DNA methylation drift

**Table 1.**
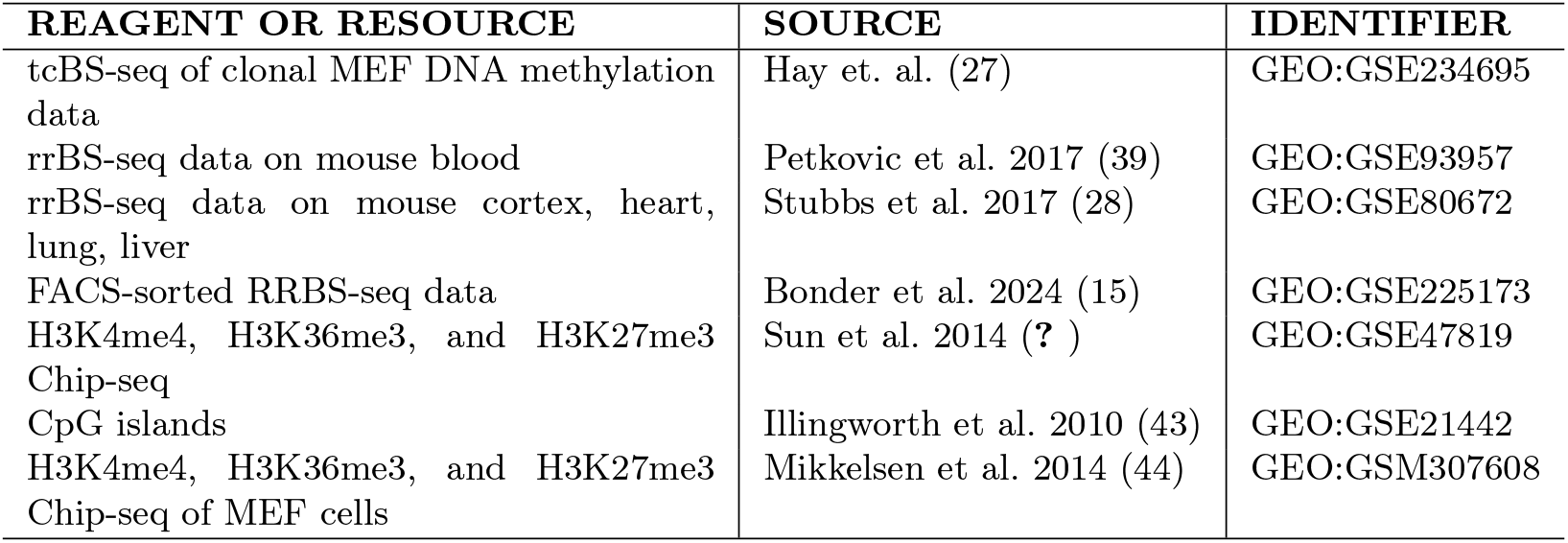
Source table.

